# Automatic Landmark Detection of Human Back Surface from Depth Images via Deep Learning

**DOI:** 10.1101/2021.02.04.429842

**Authors:** Masumeh Delgarmi, Hamed Heravi, Ali Rahimpour Jounghani, Abdullah Shahrezaie, Afshin Ebrahimi, Mousa Shamsi

## Abstract

Studying human postural structure is one of the challenging issues among scholars and physicians. The spine is known as the central axis of the body, and due to various genetic and environmental reasons, it could suffer from deformities that cause physical dysfunction and correspondingly reduce people’s quality of life. Radiography is the most common method for detecting these deformities and requires monitoring and follow-up until full treatment; however, it frequently exposes the patient to X-rays and ionization and as a result, cancer risk is increased in the patient and could be highly dangerous for children or pregnant women. To prevent this, several solutions have been proposed using topographic data analysis of the human back surface. The purpose of this research is to provide an entirely safe and non-invasive method to examine the spiral structure and its deformities. Hence, it is attempted to find the exact location of anatomical landmarks on the human back surface, which provides useful and practical information about the status of the human postural structure to the physician.

In this study, using Microsoft Kinect sensor, the depth images from the human back surface of 105 people were recorded and, our proposed approach - Deep convolution neural network-was used as a model to estimate the location of anatomical landmarks. In network architecture, two learning processes, including landmark position and affinity between the two associated landmarks, are successively performed in two separate branches. This is a bottom-up approach; thus, the runtime complexity is considerably reduced, and then the resulting anatomical points are evaluated concerning manual landmarks marked by the operator as the benchmark. Our results showed that 86.9% of PDJ and 80% of PCK. According to the results, this study was more effective than other methods with more than thousands of training data.

## 1 Introduction

Spinal deformities have always been one of the most critical issues in the medical community; the study has been dramatically increased over the past decade. Due to its particular complexities, this disease is a different and serious one and affects various age groups [1]. This disorder reduces the patients’ quality of life and is associated with pain and fatigue. Spinal deformities include Scoliosis, sagittal plane misalignment, kyphosis, slipping, and vertebrae [2]. Various factors can cause these deformities, including degenerative changes in the intervertebral discs, trauma, tumors, and infections, the occurrence of which in adulthood might be due to the development of deformities and disorders from childhood [3]. This disease’s primary issue is its early diagnosis to prevent significant clinical consequences because spinal alignment plays a critical role in protecting the nervous system and skeletal stability and maintaining the natural body alignment [1], [4]–[7].

Various invasive and non-invasive measurement systems have assessed these disorders over the decades. Radiography is one of the essential invasive diagnostic methods used for clinical purposes and is dangerous for many patients, especially adolescents, who are frequently exposed to X-rays, and its consequences may occur in the upcoming decades. Studies have shown that adolescent girls with Scoliosis are at serious risk of breast cancer [2], [3]. This could also cause breaking intermolecular forces and consequently damage deoxyribonucleic acid (DNA). Therefore, individuals during reproductive ages are prone to transfer this damaged DNA to the child via mutation. However, this phenomenon has not been proved yet [8]. Therefore, such risks have dramatically increased the necessity of using non-invasive methods for describing the spinal deformities [2].

Surface topography is the study of the 3D human back shape. This technique is an alternative for radiography to reduce the patient’s exposure to ionizing radiation during the treatment period. One of these systems’ functions that separates them from other technologies is putting anatomical landmarks on specific points of the human back surface, such as the posterior superior iliac spine (PSIS), 7th Cervical Vertebra(C7). Research has shown that this system can detect landmarks more accurately than an experienced physician [3]. These landmarks are essential for advanced topographic analysis because they are regarded as a fixed coordinate system and an axis for the patient [4].

In this study, a new heuristic approach automatically detects anatomical landmarks of the human back surface. The system includes a Kinect depth sensor [9]. The resulting information is a grayscale image of the human body surface that contains information about the exact location of critical points. Given that recent approaches based on deep learning have led to significant achievements in various machine vision studies [10], thus, in this study, the problem can be examined using deep learning methods and convolutional neural networks (CNNs). It should be noted that the methods are utilized to detect the critical points of the body in the human pose estimation because restoring the human condition means locating the critical points of the body. Although this research aims not to estimate the body pose, the anatomical landmark points have a specific description of it.

Deep learning methods also pose new challenges, the first of which is the lack of training images due to the limitations of recording images from countless people. To compensate for this problem, the data augmentation method was utilized. The second challenge was to train an optimal model to understand and describe the human back surface features from depth data. Training a Neck-network from scratch is not a good idea, especially when training it by a small dataset. We used a pre-trained model as a feature descriptor for other parts of our model to cover this. Therefore, the transfer-learning technique was applied in network architecture by using ResNet-152. In this research, the network was separately implemented with three feature descriptors, including ResNet-50[11], VGG-19, ResNet-152[12], and its performance was compared in all three models. We predict that Resnet-152 could be the best choice to describe the depth data features.

## 2 Related Work

### 2-1 Landmark Detection without Deep Learning

The nature of the data we used plays a significant role in selecting the appropriate method for the automatic analysis of the human back shape. These are application-dependent methods; thus, any methods that have been made so far must be consistent with the data used [1]. In this research, the depth data were used and approximated as (x,y), indicating each landmark’s coordinates on the image. It should be noted that there are several methods based on 3D segmentation. L. Soler et al. [13] used this method to analyze anatomical structures of 3D volumetric data obtained from CT. This method is not useful for 2D data, in which the structures are placed on a surface. Some other methods attempt to describe the anatomical shape feature achieved by marking landmarks on the body surface. For example, M. A. Styner et al. [14] used the spherical harmonics (SPHARM) method, which provides a significant representation of the surface for shape analysis. This method only works on data with spherical topology and could not support the data used in this research. Another method is the active shape model (ASM) [15], a statistical model-based feature matching method used to detect the object’s shape in an image using the point distribution model for statistical matching. In this method, the shapes are described by a set of points; these points are allowed to deform following the original deformation. This method was initially developed for working with 2D images having the desired embossing. The point distribution model (PDM) can be easily generalized to be consistent with the 3D nature of input data by adding third coordinates during analysis. This is valid for volumetric data, in which the model can be entirely deformed in all three dimensions. This method does not work appropriately for detecting deformities in the body’s back structure because a small percentage of the total population has a deformation problem on the human body surface. Thus, in the ASM method, a very coherent training set that does not contain all changes cannot find new shapes that are valid in realtime. Therefore, these changes can be unintentionally neglected during modeling by principle component analysis (PCA) [1], [16]. In short, although the described approaches provide an invaluable insight into data analysis, they do not correctly solve the problem due to the above limitations.

### 2-2 Landmark Detection with Deep Learning

CNNs have achieved advanced results in various fields due to being resistant and having high learning capacity [7]. The human pose estimation aims to approximate the location of the human joints in images. In other words, it is a process that involves finding key points such as arms and shoulders and then combines them in a model [17]. In this section, we mainly discuss deep learning methods to approximate these points. In general, based on the management of input images, the deep learning methods can be classified into holistic- and part-based methods [17]. DeepPose is an example of a holistic-based approach that was proposed by Toshev et al. [18]. This method is considered the first approach used in deep learning to estimate the human pose and is formulated as a regression problem, and the idea of this method in correcting and improving the estimations is to use the cascade of regressors [19]–[21]. This model has made progress in several challenging datasets and attracted researchers’ attention to CNNs. The holistic-based models are not very accurate because it is challenging to learn the regression of point vectors directly from images. In the part-based method, the body parts are approximated and combined by a graphical model [17]. Thomson et al. [22] used a combination of CNNs and graphical models to estimate components. The graphical models often learn the spatial relationships and distances between the points. In this method, the network output is heatmap points, instead of continuous regression, estimating a landmark’s probability in each pixel. The results of this method are very successful and lead future works towards heatmaps. This method’s only drawback is the lack of body structure modeling and not detecting the point-to-point connection. Establishing this affinity is very important from the viewpoint of the detected vital points because the formation of this affinity leads to the ease of displaying the key points. Therefore, this problem was solved by the following methods. Wei et al. [23] proposed a completely different method to estimate multi-stage CNN-based human pose, and a part of the body is estimated at each stage. Therefore, fewer errors could occur in landmark detection. This network uses an intermediate layer after each stage due to errors in dealing with decreasing slope during training. Carreira et al. [24] proposed an algorithm known as duplicate error feedback. Using predictive error feedback, the designed model corrects its initial prediction several times. This method can be regarded as estimating an approximate mode to estimate a more accurate mode. Cao et al.[25] extracted features from the first ten layers of VGG-19[11]. Then, these features were processed using a three-output CNN with two input sources, and one output predicts the location of critical points. One output retains the vector field maps, and the third output retains the initial features. This method is performed iteratively. Multi-stage CNN combines the processing results of each of CNN’s outputs, which is used to obtain rich information from images to improve the function [25]–[28]. Hence, given that this algorithm is the most advanced method in this field and has multiple outputs, this method can detect the desired landmarks for medical purposes. On the other side, having three outputs is one of the advantages of this method. In detecting anatomical landmarks on the human back surface, by determining affinity between two pairs of the related points, a view of the human spine can also be predicted. Despite all the advantages of this method, after testing and implementing the network proposed by Cao et al. [25], the result showed that VGG-19 [11] is not appropriate as a feature descriptor for our depth data, and no good output is obtained. Thus, in this paper, the structure of the feature descriptor was changed and Resnet-152 was employed. The results show that the used algorithm achieved the best results even despite the lack of training data. In the following, we will elaborate on the algorithm operation. In Section 3, we indicate show how to choose the anatomical structure and network architecture. Section 4 describes how to collect data and Section 5 explains the preprocessing and data enhancement methods. Section 6 shows the evaluation criteria and Section 7 describes the network training along with the network parameters. We will compare the results of applying the algorithm to the VGG-19, Resnet-50 and Resnet-152 networks, and finally both discussion and conclusion are provided in Section 8.

## 3 Methodology

This approach’s primary purpose is the automatic landmark detection of the human body surface and the affinity to evaluate the spinal deformities and ease of the diagnostic process. As mentioned, the restriction of traditional methods and shared goals with human pose estimation provides the context to study the latest methods in the literature. Thus, in this research, human pose estimation’s best architecture is inspired for medical purposes. In this section, first, choosing the key points related to spinal displacements, and then, the deep convolutional network architecture used for the stated purposes is described in detail.

### 3-1 Selection of Anatomical Structures

A topographical assessment of the human back surface visually provides the physician’s postural structure’s required information. However, this requires determining a series of variables and indicators to detect disorders and deformities of the human back surface using these indicators. Two indicators that have been widely used as determinants to this date are called posterior trunk symmetry index [29] and deformity in the axial plane index [30]. The location of the landmarks is optionally defined. A common feature of these structures is that their position is influenced by the shape of the spine that can facilitate the detection of deformities using topographic data [1].

This paper is inspired by the list of landmarks in both deformity in the axial plane index (DAPI) and posterior trunk symmetry index (POTSI) indicators, and concerning the definition of the problem, 10 points are shown in Figure 1 were selected as anatomical structures. These structures contain a small area and no single points. This means that these points’ neighborhood can be considered an anatomical landmark, but the parameters used to evaluate the deformities should consider single positions as an input. The output of the described algorithm must also be single-point, each point of which indicates the position of an anatomical structure [1], [2].

**Figure 1:**
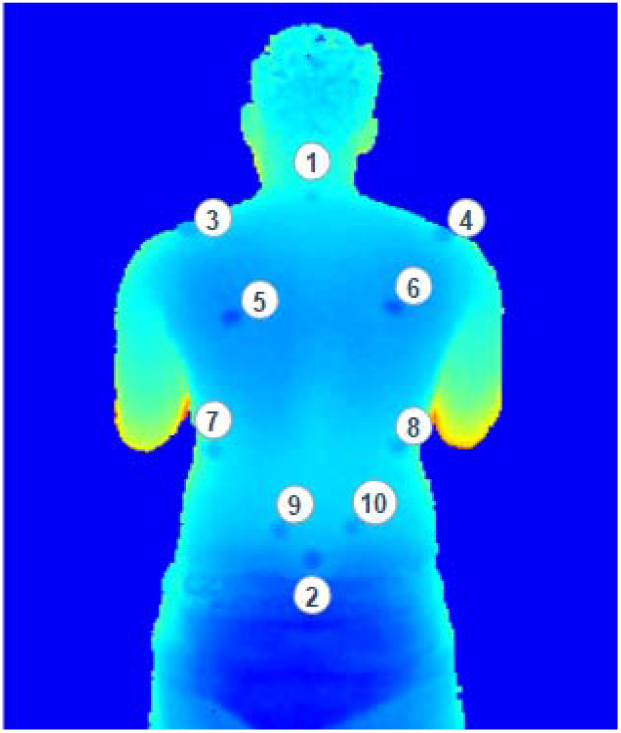
Illustration of 10 anatomical landmarks.

### 3-2 Model Architecture

The proposed network architecture consists of two parts. 1. Feature extraction, 2. Prediction of landmark position and the affinity. In the present research, the data’s characteristics have caused the complexity of feature extraction because understanding the depth data features for medical purposes is highly sensitive. Then, choosing an appropriate feature extractor plays a determinant role in the network’s accuracy and performance. In this study, ResNet-152 [12] is selected as a feature descriptor. It is noteworthy to mention that to match the input image to the standard structure of the network ResNet-152, a stacked image of the input was created and applied to the network input.

The output of this stage is fed to the n stage. As illustrated in figure 2, each stage is divided into two parts. The first branch is made of a deep convolutional network that produces a heatmap for each landmark and is defined as *H* = (*H*_1_, ...., *H*_*j*_). The design of branch 2 is the same as branch 1, except that the network in this branch tends to predict the affinity between these points, which is known as landmark affinity field and is defined as *L* = (*L*_1_, ... ., *L*_*C*_). There is a vector for each joint between the two landmarks. Each pixel on the heatmap indicates the possibility of a specific key point. These two branches are considered a stage in this architecture, and as required, stage 7 is used in this work [25]. The network architecture is shown in Figure 2.

**Figure 2:**
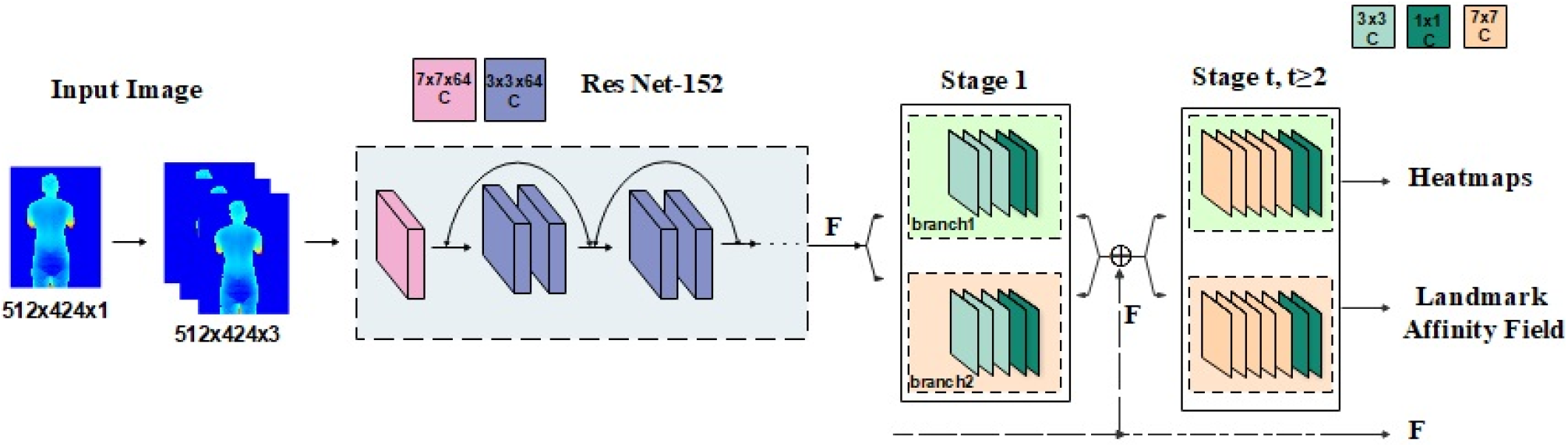
Architecture of the two-branch multi-stage CNN.

## 4 Dataset Collection

In this study, data were collected by Microsoft’s Kinect-v2 [9]. Fifty women and fifty-five men aged 19-35 years old were recruited at Tabriz University of Medical Sciences, which resulted in 105 depth images of the human back surface. The participants were assumed to be healthy and have no history of Scoliosis. These people were asked to naturally stand with their backs to the camera so that the distance of feet from the vertical line was identical (See an example in figure 3). The Kinect sensor was located 1.5 m behind the participants. As shown in Figure 3, the data entry process was as follows: at first, a marker was placed at point C7, and imaging was done without other landmarks. Then, the rest of the markers were embedded in the exact location, and the second image was recorded with landmarks. The images were saved as raw images in the size of 512×424. It is necessary to note that due to the presence of error sources in the Kinect sensor, we decided to limit the image conditions to a closed area far from direct light, and a 20-min delay was required to put the sensor in stable conditions [31]. The Institutional Review Board approved the protocol for research ethics and human subjects protection at Tabriz University of Medical Sciences. All participants gave informed consent after the experimental procedures were explained to them.

**Figure3:**
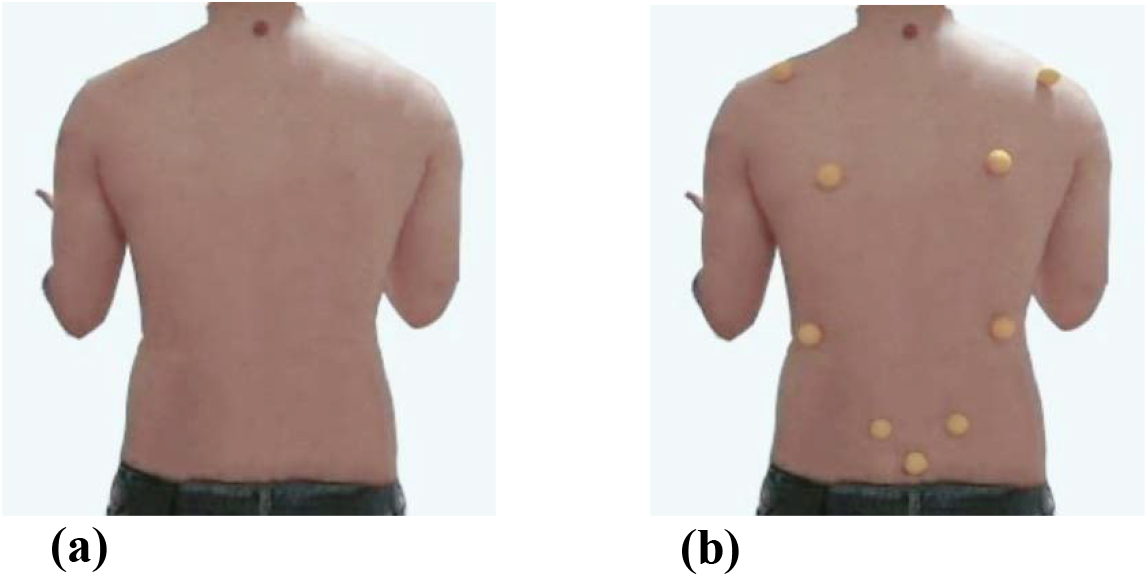
Example of captured images

## 5 Preprocess and Augmentation Pipeline

Given the characteristics of the Kinect sensor, the resulting depth images were very noisy [9]. Hence, it was necessary to apply a pre-processing stage to improve the quality of the depth image. Thus, if r were the depth image matrix, it would be removed with the thresholding method *r* > 1.7*m*. Figure 4 shows the depth image before and after noise removal.

**Figure 4:**
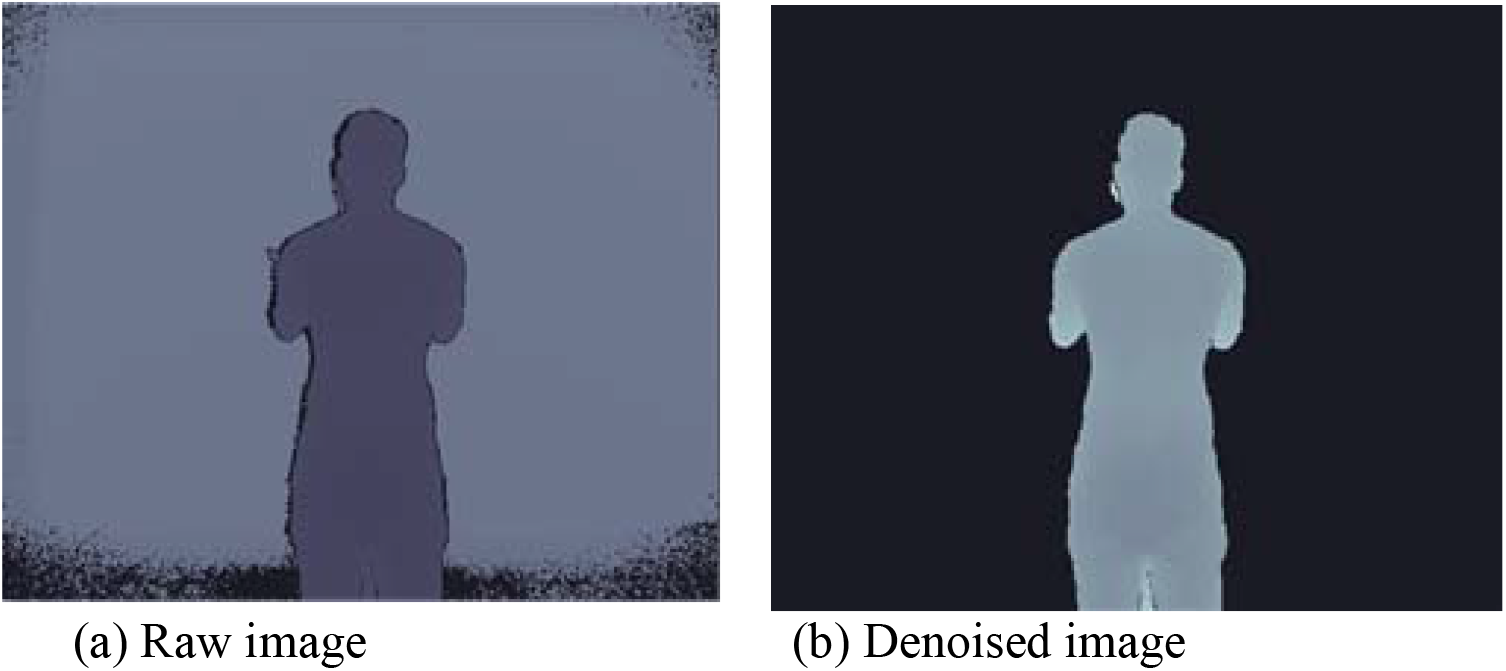
Example of Raw image and Denoised image

In the depth images, the range of pixels varied in different scenes. Meanwhile, the sensor error and movement of objects in the scene dynamically changed the minimum and maximum distance. Thus, it made feature extraction more difficult during convolutional operations. Therefore, in the training process of CNNs, the network was hardly converged. To solve this problem, the data was normalized through equation 1. Therefore, the data range was converted to a range of 0 and 1[32].

In Equation 1, R is the input image matrix and is the post-normalization raw image.

The first step in using CNNs is to have an extensive dataset. In fact, these networks have provided a platform that needs a large volume of data to train the network without overfitting. If we want to use deep learning for specific expertise, this need still exists. Compared with large datasets used in deep learning, such as COCO [33] and ImageNet [34], the dataset collected for such research, which requires precise detection and high sensitivity, is very small. Then, data augmentation [35] was used to solve the data shortage problem. We applied four transfer techniques to the right, left, top, and bottom of the dataset’s images. Figure 5 shows an example of the images. The final number of images of the dataset was 525.

**Figure 5:**
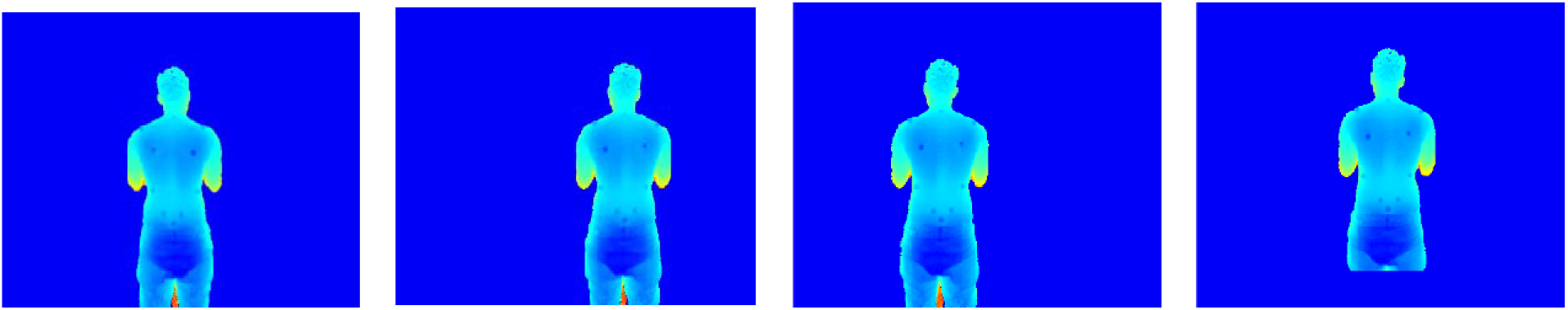
Example of Data Augmentation

## 6 Evaluation Metrics

In automatic landmark detection, two well-known criteria of PCK (i.e., the percentage of points that are correctly estimated) [36] and PDJ (i.e., the percentage of correctly detected points) [37] are widely used. First, PDJ is the percentage of points that are correctly detected. A point is detected correctly when the distance between the predicted location and real location is less than a coefficient of the upper body’s diameter. The upper body’s diameter is defined as the distance between the left shoulder and the right thigh. This coefficient is considered equal to 0.2. Second, PCK is the percentage of points that are correctly estimated. A point is correctly estimated when the distance between the predicted location and real location is less than a threshold. This threshold limit can be considered equal to 50% of head length, less than 0.2 of upper body diameter, or equal to 150 mm.

## 7 Experiment and Results

### 7-1 Training Dataset Generation

After recording the data, the training dataset’s labeling process should be performed immediately for network training [38-40]. As mentioned earlier, the network has a branch for learning the position and spatial coordinates of landmarks and another branch for finding affinity between the two related points. Thus, all the information relevant to the JSON file’s training images was set in the defined format in the first step. This information included the saved image address, image size, spatial coordinates of landmarks, and ID of each image, and the connection points. For example, “Landmarks” represents an image’s landmark coordinates from a dataset written as (x, y) from 1 to 10 in a row. “Links” also represents the number of both connection points within a bracket, e.g., points 9 and 10 are connected.

“Landmarks”: [267,137,261,304,214,154,313,154,239,202,294,192,227,261,295,258,247,286,273,291]

“links”: {[[9,10],[7,8],[5,6],[3,4],[1,2]]},...]}

Using this information, heatmaps and vector field maps for each image are labeled for network training. In the next section, we will explain how to create them with the existing mathematical equations.

### 7-2 Heatmap Generation

For this issue, a label is made for each landmark as groundtruth using the Gaussian distribution, the mean of which represents the location of landmarks in the image. Each heatmap indicates the position where the landmarks occur. These heatmaps are derived from the coordinates of the critical points stored in the JSON file.

Assume 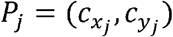 is the correct position of the j^th^ landmark for the person in the image so that *x*_*j*_, *y*_*j*_ ∈ *R*^2^. To make a heatmap for this landmark, it is necessary to define a neighborhood to the center of this point because it is determined that each heatmap does not contain only one point but also involves a small area, and then the value of the Gaussian function for *P* = (*x, y*) is calculated in this area. A heatmap for the j^th^ landmark is shown with 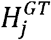.

Then, the value of the Gaussian function in *P ∈ R*^2^ position for the j^th^ landmark is defined as follows [14]:

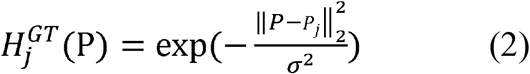

Equation 2 shows the 2D Gaussian function at the center of the correct landmark position. An example of a heatmap resulting from this method was illustrated in figure 6.

**Figure 6:**
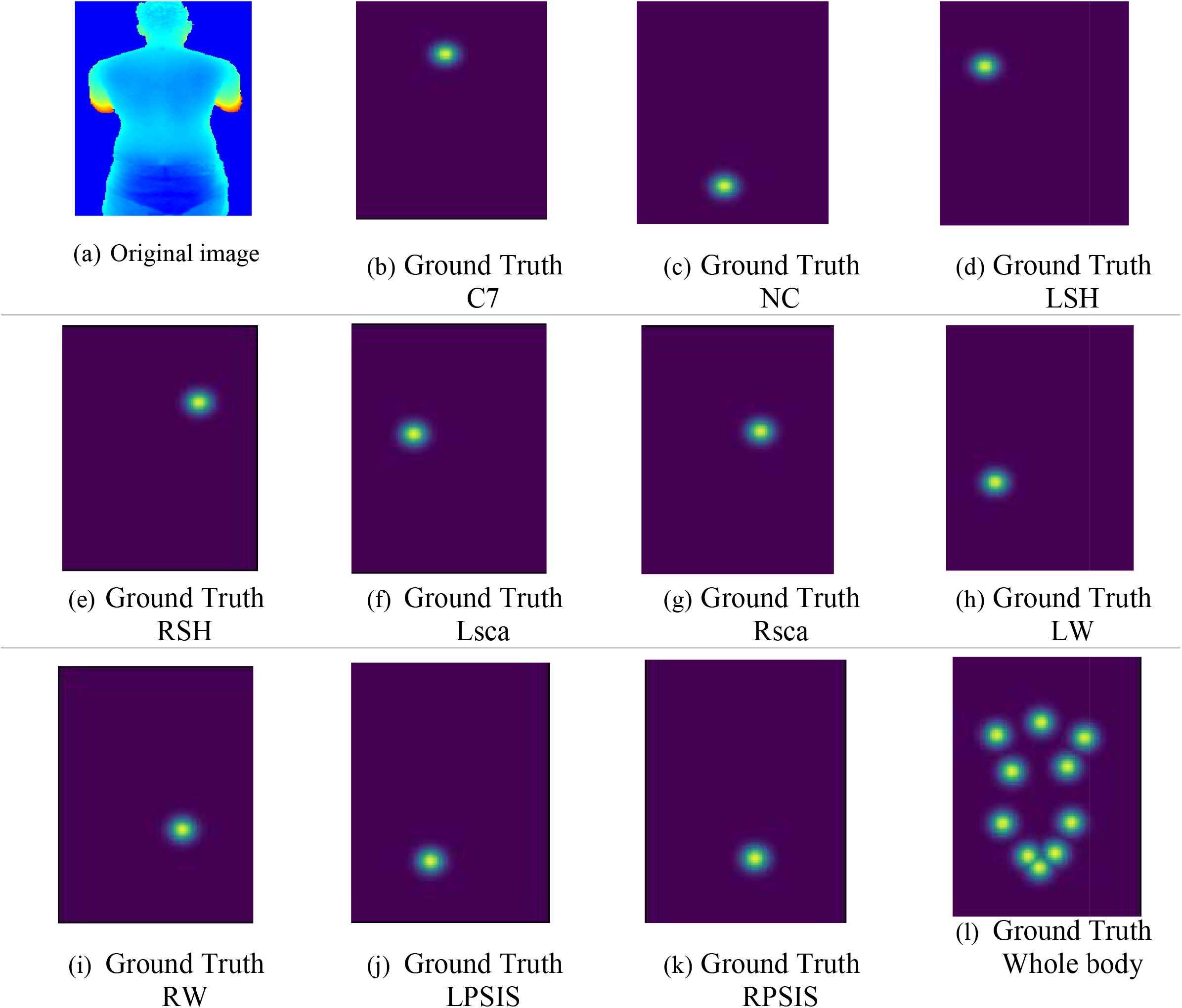
Example of Heatmaps

Below, we indicate how to formulate and generate ground truth images for anatomical landmarks mathematically. As mentioned, this method is based on the Gaussian function. First, an affinity around *c*_*x*_, *c*_*y*_ is defined, and the value of the Gaussian function corresponding to all x and y values in this affinity is computed. We also show how to generate groundtruth images for automatic landmarks

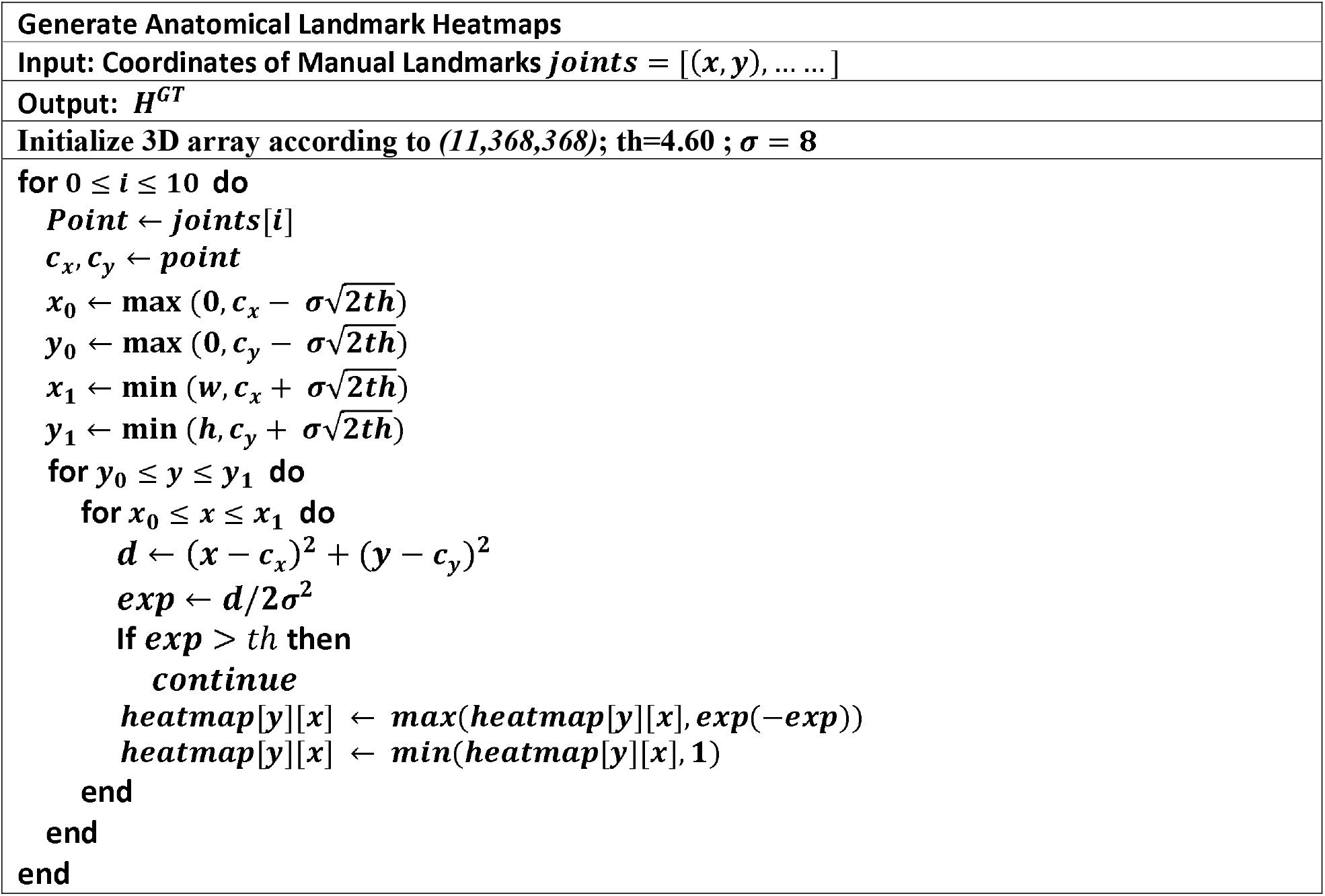

### 7-2 Landmark Affinity Fields Generation

Each of the defined joints holds a vector map that connects two different parts of the body and contains the position and direction information in the joints. In Figure 7, a joint is shown between two landmarks. Assume that and are positions of the two related landmarks and in the image. If point is on the joint c, the value of is a single vector; otherwise, a vector with zero value for all the other points [19].

**Figure 7:**
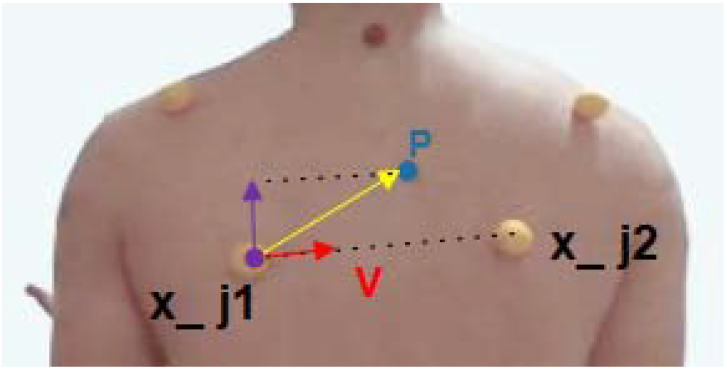
Landmark Affinity Field

A vector filed map in the image, points p is defined as follows [14]:

(3)

where v is a single vector in the direction of joints and is defined as follows:

(4)

Figure 8 shows an example of vector maps obtained by the above method.

**Figure 8:**
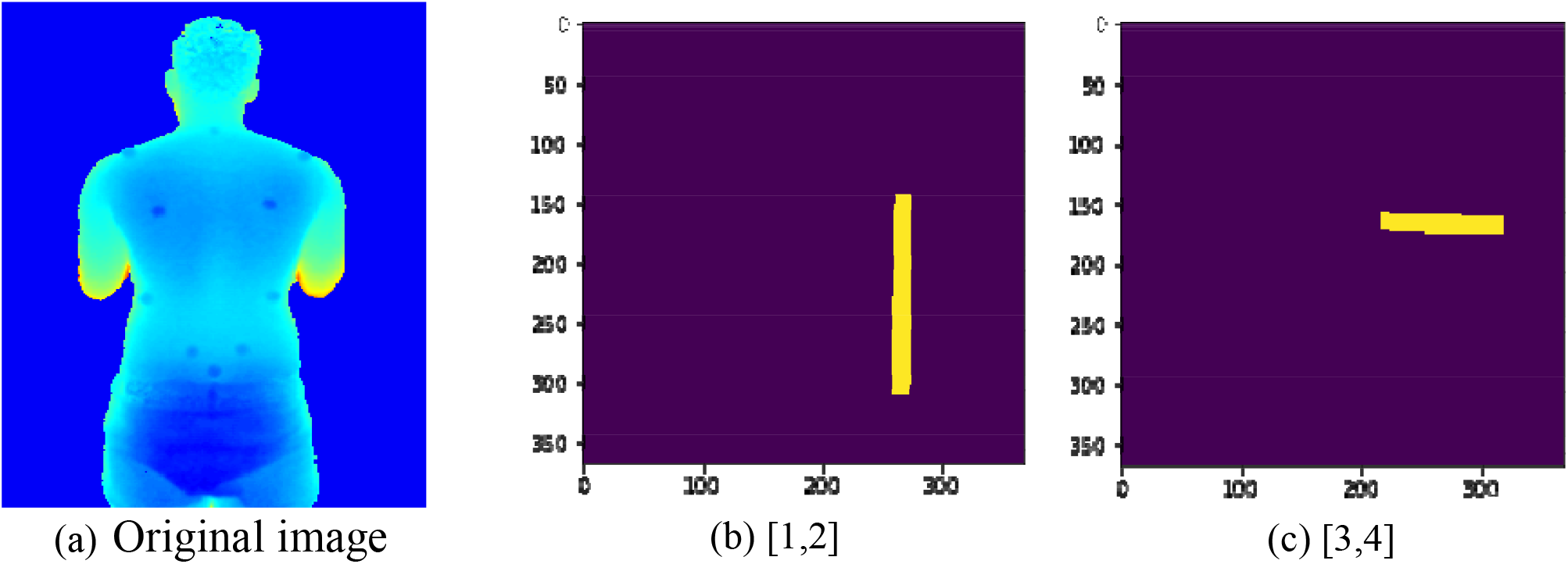
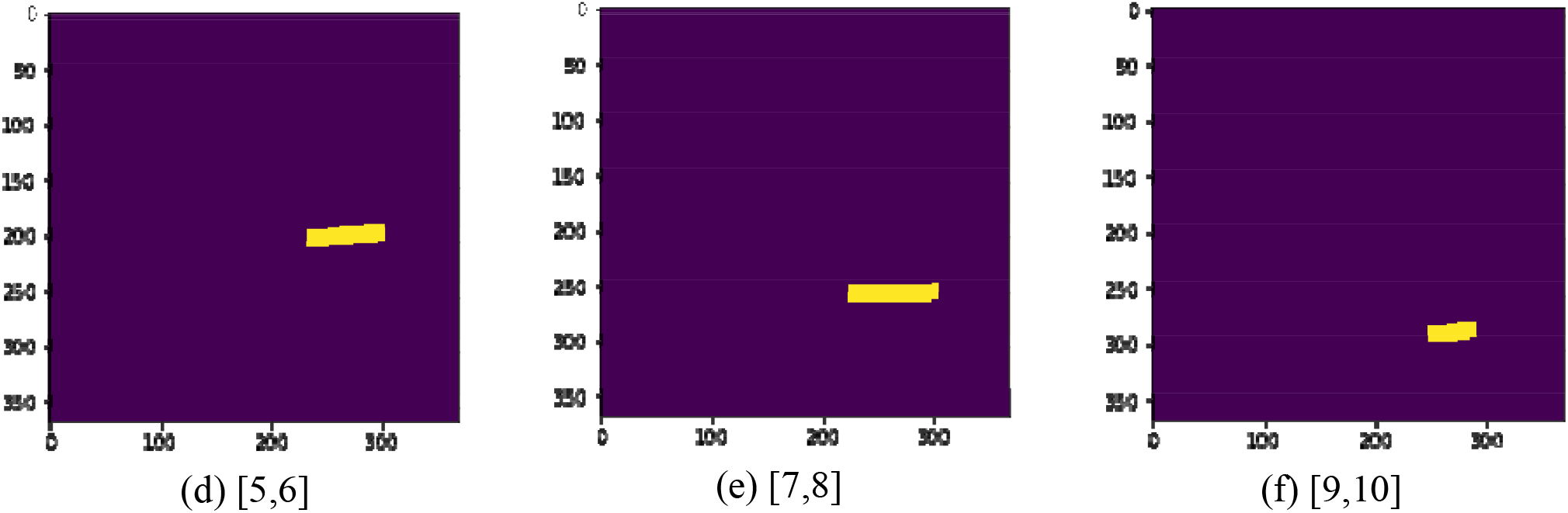
Example of vector map

### 7-3 Training Protocol

We trained the model on 400 images and used the rest of the images for network testing. The deep learning model is created in Python 3.6 and TensorFlow 1.4. To design an optimal model, the heatmaps and vector maps were trained simultaneously using a training process similar to the training process [19]. Our model’s best performance was achieved with a batch size of 3 and a learning rate of 0.0001. It should be noted that maximum epochs were equal to 70, and the network was implemented on a system with Nvidia Tesla T4 and 12GB RAM.

### 7-4 Training

Figure 2 shows that the image (424 × 512) is entirely imported into the convolutional network Resnet-152. Transfer learning occurs in this part because the network extracts the features of the input image without supervision. The result is a set of feature maps F that are entered into each branch, and local features are obtained. In this stage, the layers are more meaningful because they are directly set with the training data label.

Branch 1 predicts anatomical points, and Branch 2 can detect the affinity between two pairs of related points and show them as vector maps at the output. This part is significant since the deformities can be visually and generally found by drawing these affinities in the human back front structure, which requires a specialist to detect the affinity between joints with various spinal diseases.

Assume that and are the same CNNs. Then, the first branch produces a set of heatmaps and the second branch produces a set of vector map. In the next stages, both branches’ predictions in the previous stage and features of main image F obtained from the descriptor are combined.

Equations 5 and 6 show the predictions of branches 1 and 2 in step t, respectively [19].

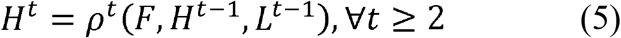

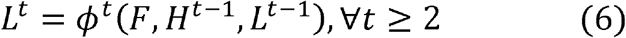

To guide the network and correct predictions, two-loss functions are applied at the end of each step, where 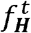 and 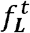 are used for branches 1 and 2, respectively. The specific cost function is assigned to each branch per step.

Loss functions in both branches in step t are defined as follows:

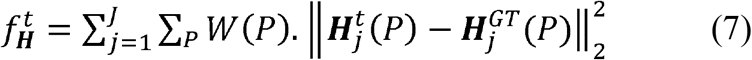

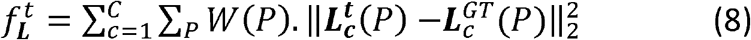

where 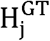 is the ground truth of heatmap, 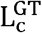 is the ground truth of vector map, and W is a binary mask with *W*(*P*) = 0. This binary mask made of ground truth is applied to prevent the removal of correct predictions to the outputs to mask the points. P represents a pixel of the image. Equation 9 shows the total cost function, which is a combination of the cost functions 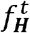 and 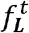.

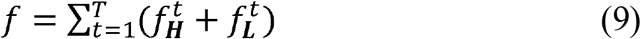

## 8 Results

### 8-1 Results on VGG-19

First, we were required to design and implement a descriptor to comprehend and extract the general concepts and input distinguishing features. This becomes problematic because of having depth images with different visual content compared to the color images. Putting the model under training from scratch because of leakage in our depth images dataset was not possible. Therefore, we tried to see transfer-learning from pre-trained models over color images to work on depth images. First, the deep convolutional network VGG-19 is utilized as a feature extractor, but no satisfactory results are achieved.

Figure 9 depicted a 10-channel Heatmap. Each of these heatmaps is the output of the first branch. As shown, the predicted Heatmaps are expected to represent anatomical landmarks of the human back surface. However, the desired output is failed to be obtained. Figure 10 also shows the Affinity part map. The undesired network performance in detecting and extracting the distinguishing features in this algorithm is observed.

**Figure 9:**
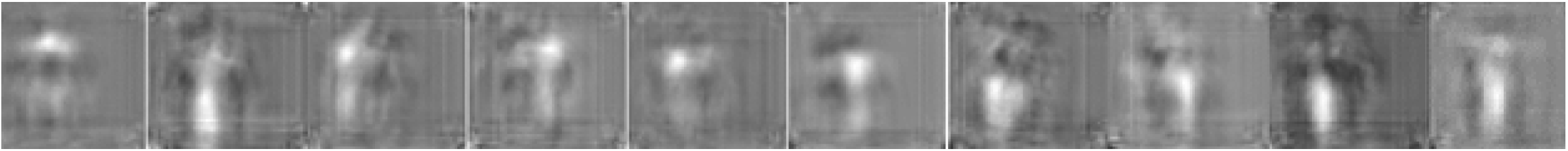
Confidence map of stage-7 (trained with VGG-19)

**Figure 10:**
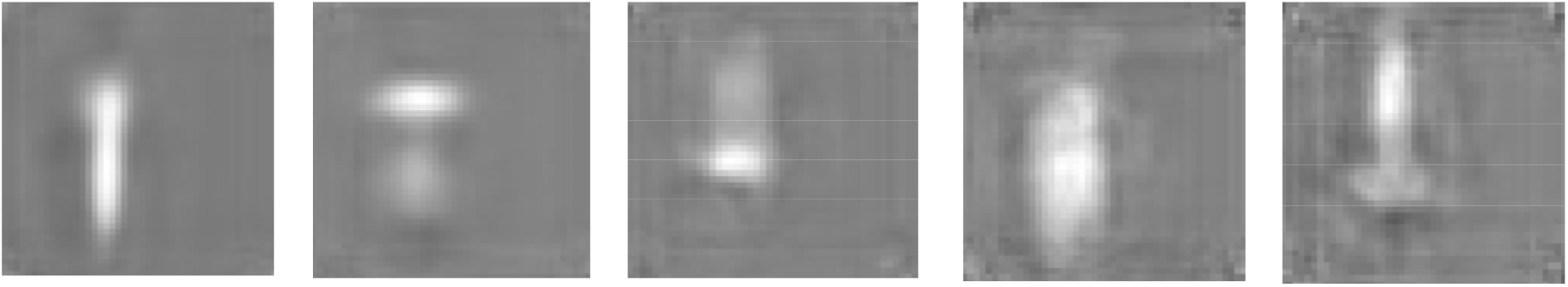
Vector map of stage-7 (trained with VGG-19)

### 8-2 Results on ResNet-152

Regarding the poor function of the VGG-19 network, the ResNet-152 network was applied. The network was expected to have better performance than VGG-19. Figure 11 shows the prediction of heatmaps obtained from the last layer of the CNN, and Figure 12 represents the prediction of vector maps.

**Figure11:**
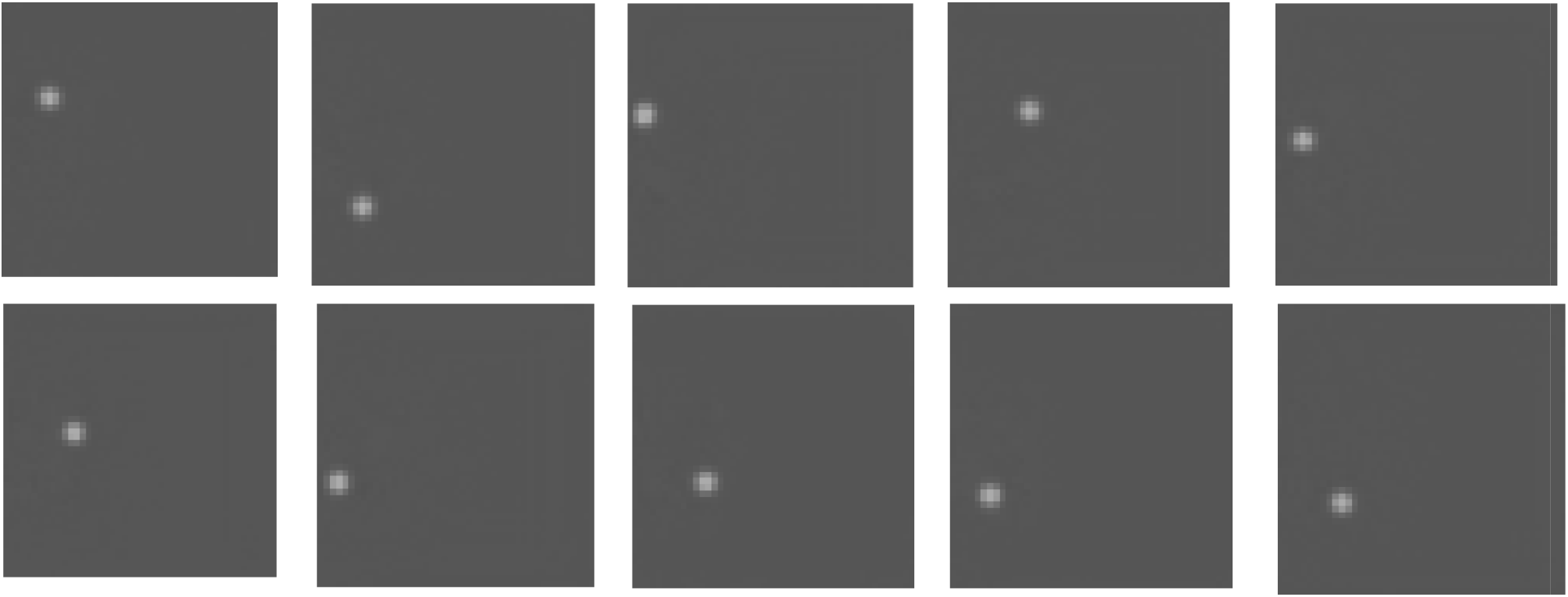
Confidence map of stage-7 (trained with Resnet-152)

**Figure12:**
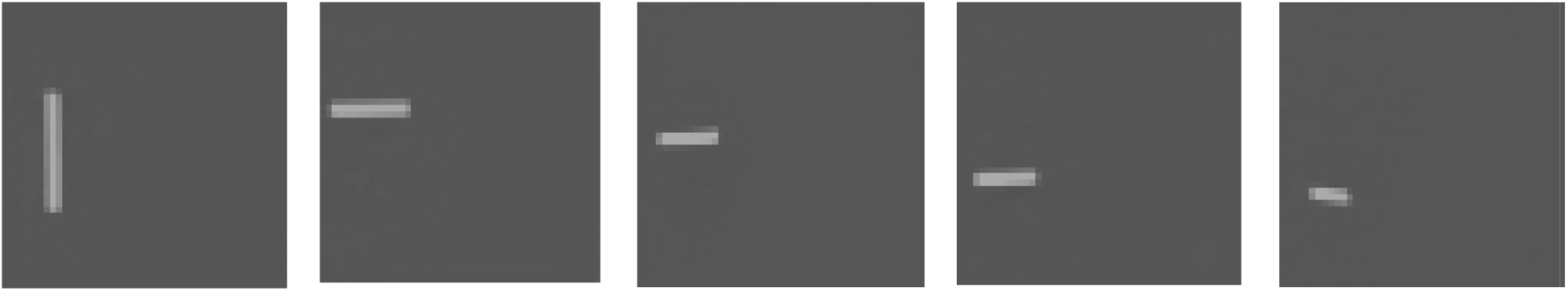
Vector map of stage-7 (trained with Resnet-152)

By applying the input image to the ResNet-152 descriptor, the network could comprehend and extract distinguishing features from the input image. The second part of the network architecture could quickly learn the extracted features and achieve remarkable and promising results. After the training process ended, the experimental data were used to test the new architecture and ensure proper training.

In addition to increasing network accuracy, ResNet-152 can be used as a feature descriptor to accelerate the network convergence process. This result was expected since this network was the deepest version of the ResNet network and had higher accuracy than ResNet-50 in the ranking. Therefore, this network’s outputs, which will be presented in more details in the next section, obtained higher percentage in proper landmark detection compared to others based on the evaluation criteria.

In Figure 13, the results of applying the evaluation samples to the ResNet-152 network are shown, which was visually compared with the ground truth of each one, and the process of improvement can be observed in network performance. The red points and lines indicate ground truth in these images, and the yellow points and lines indicate the predicted output. Here the network had a very remarkable performance in detecting all points, although the participants were assumed to be healthy. However, in practice, there were cases with spinal deformities and Scoliosis, an example of which can b observed in Figure 13. In addition to automatic landmark detection, the proposed algorithm could predict a view of people’s spine by finding a relationship in these points; this will help diagnose Scoliosis and other spinal deformities.

**Figure 13.**
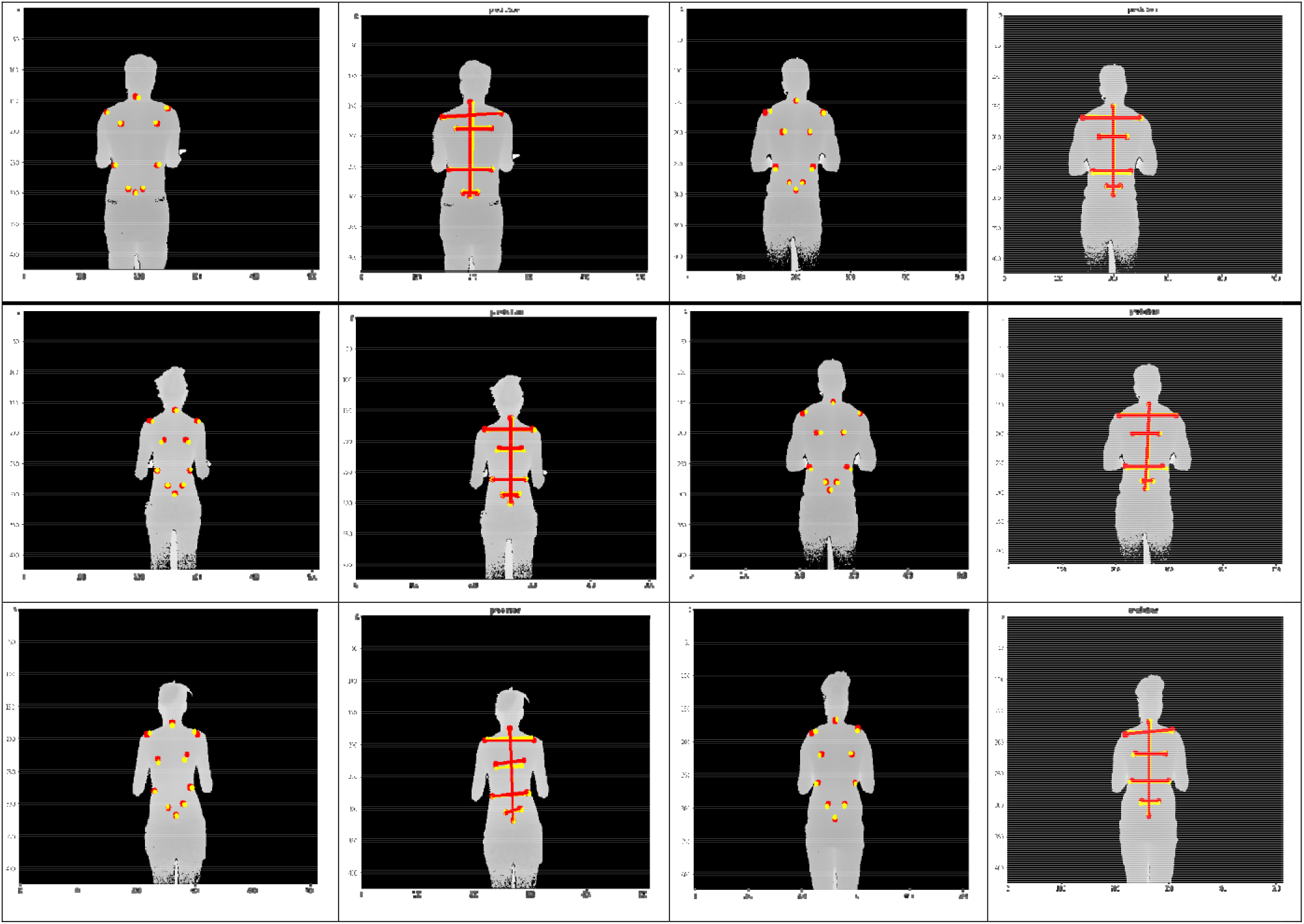
Sample plots of output from our model

**Figure 14:**
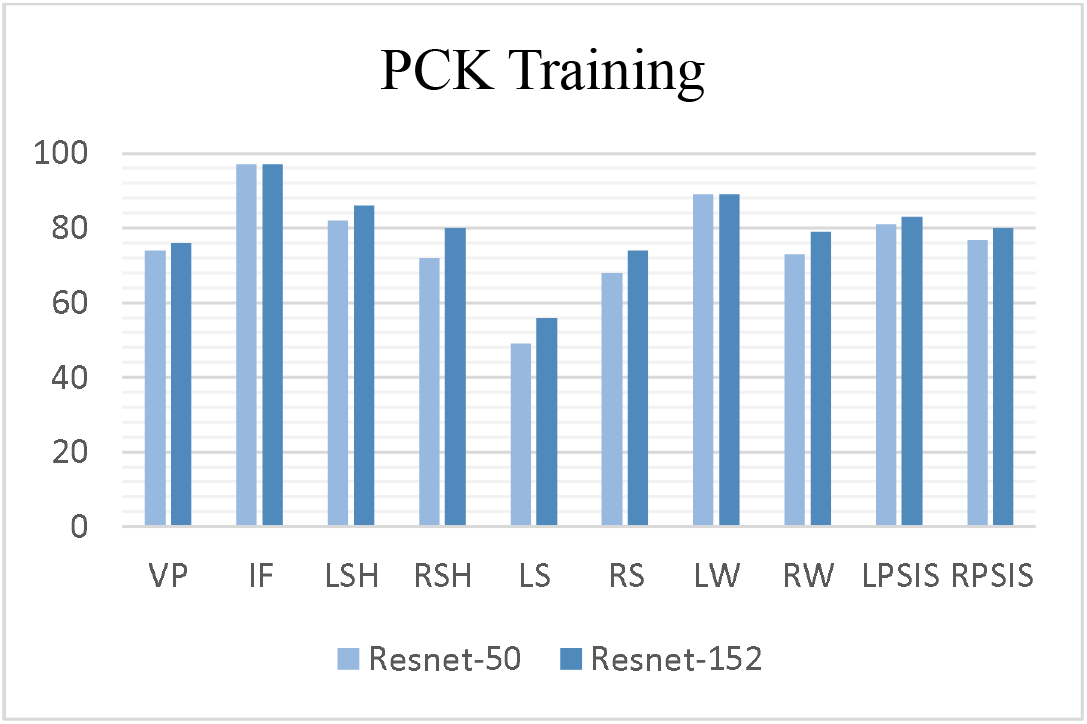
PCK comparison in training phase between Resnet-50 and Resnet-152

**Figure 15:**
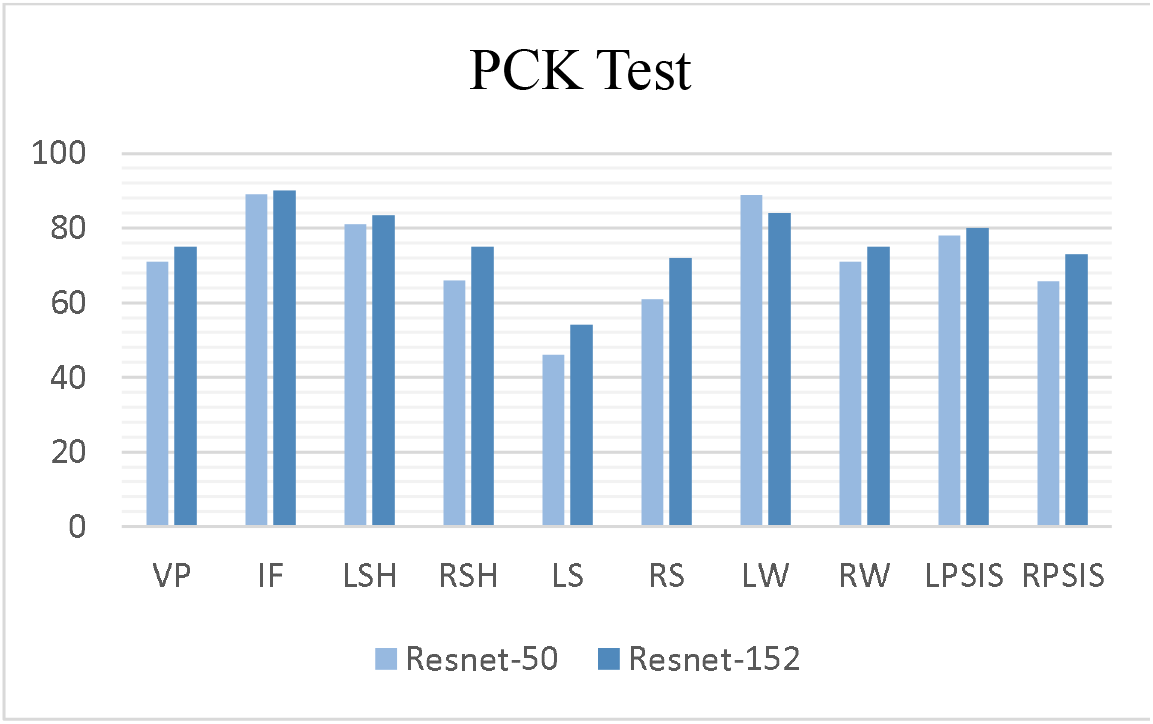
PCK comparison in testing phase between Resnet-50 and Resnet-152

**Figure 16:**
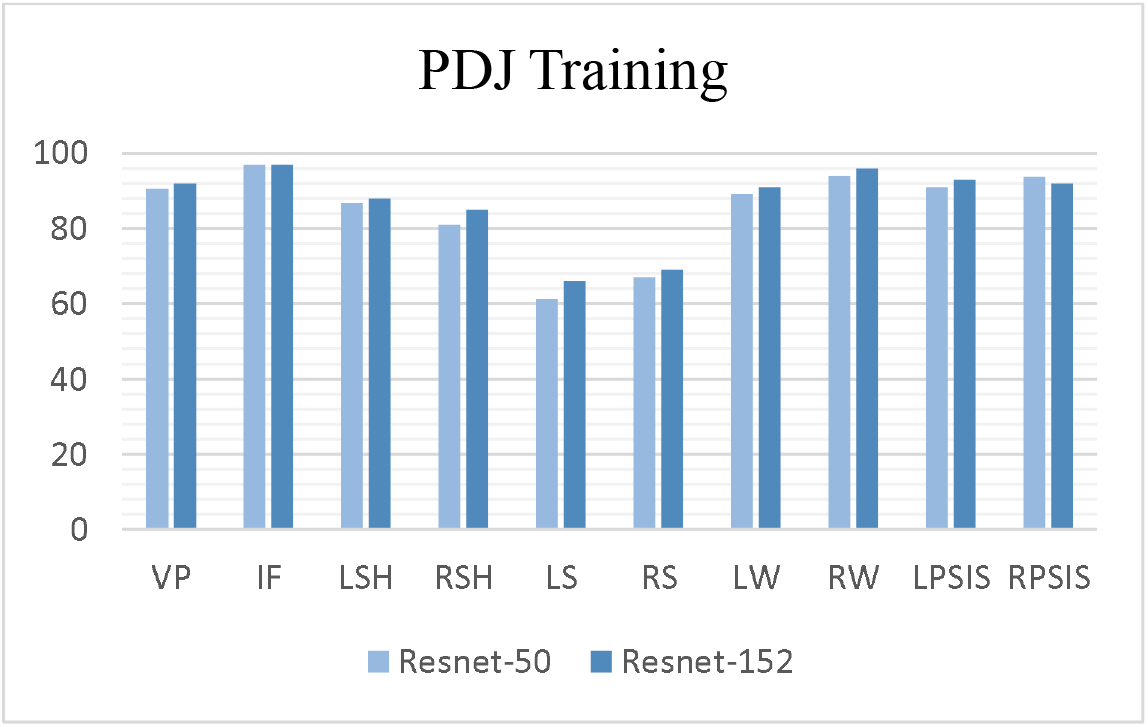
PDJ comparison in training phase between Resnet-50 and Resnet-152

**Figure 17:**
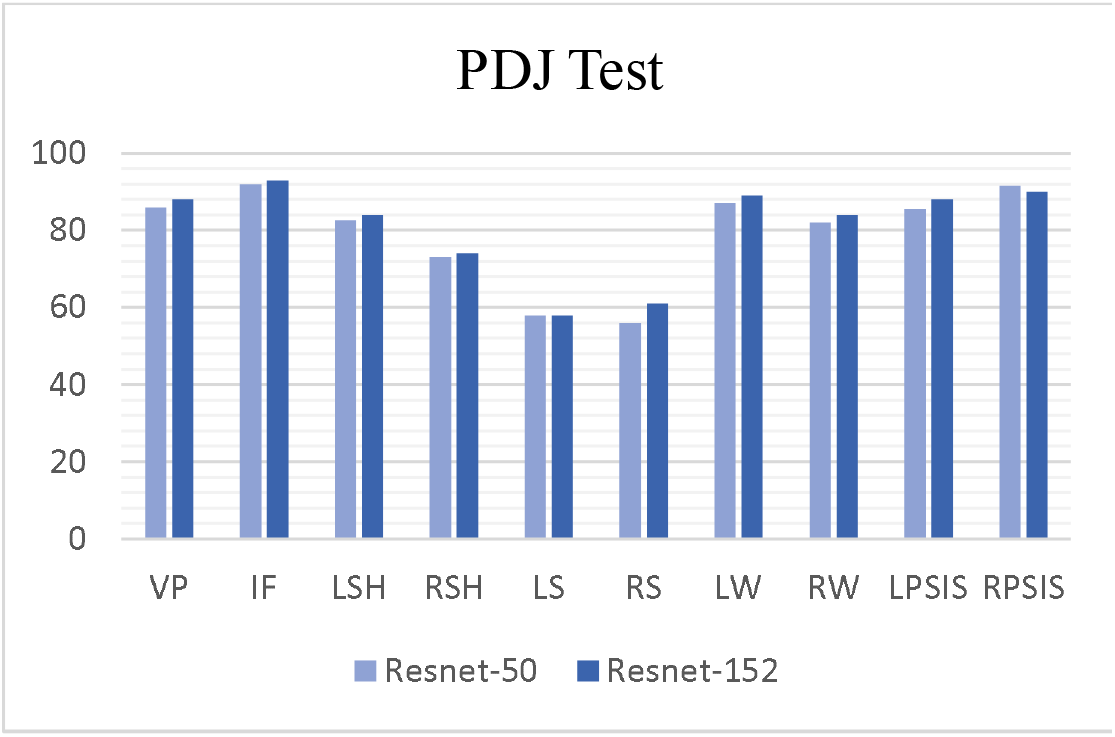
PDJ comparison in testing phase between Resnet-50 and Resnet-152

### 7-5 Comparisons

Evaluation criteria were calculated for both ResNet-50 and ResNet-152 networks. In this comparison, the VGG-19 model was not included, because it did not produce comprehensible output. By comparing both networks’ performance, as expected, the final version of ResNet increased network accuracy with less difference from the original version. Table 2 illustrates the PCK for all landmarks.

**Table1:** List of single points.

| Anatomical Landmarks |  |
| --- | --- |
| 1 | Vertebra Prominens |
| 2 | Natal Cleft |
| 3 | L Shoulder |
| 4 | R Shoulder |
| 5 | L Scapulae |
| 6 | R Scapulae |
| 7 | L Waist Line |
| 8 | R Waist Line |
| 9 | LPSIS |
| 10 | RPSIS |

**Table 2:** PCK results

| Method | VP | IF | LSH | RSH | LS | RS | LW | RW | LPSIS | RPSIS | Total |
| --- | --- | --- | --- | --- | --- | --- | --- | --- | --- | --- | --- |
| **Train** |  |  |  |  |  |  |  |  |  |  |  |
| ResNet-50 | 74 | 97 | 82 | 72 | 49 | 68 | 89 | 73 | 81 | 76.8 | 76.1 |
| ResNet-152 | **76** | **97** | **86** | **80** | **56** | **74** | **89** | **79** | **83** | **80** | **80** |
| **Test** |  |  |  |  |  |  |  |  |  |  |  |
| ResNet-50 | 71 | 89 | 81 | 66 | 46 | 60.9 | 88.8 | 71 | 78 | 65.7 | 71.7 |
| ResNet-152 | **75** | **90** | **83.4** | **75** | **54** | **72** | **84** | **75** | **80** | **73** | **76.1** |

**Table 3:** PDJ results

| Method | VP | IF | LSH | RSH | LS | RS | LW | RW | LPSIS | RPSIS | Total |
| --- | --- | --- | --- | --- | --- | --- | --- | --- | --- | --- | --- |
| **Train** |  |  |  |  |  |  |  |  |  |  |  |
| ResNet-50 | 90.6 | 97 | 86.7 | 81 | 61.2 | 67 | 89.1 | 94 | 91 | 93.6 | 85.1 |
| ResNet-152 | **92** | **97** | **88** | **85** | **66** | **69** | **91** | **96** | **93** | **92** | **86.9** |
| **Test** |  |  |  |  |  |  |  |  |  |  |  |
| ResNet-50 | 86 | 92 | 82.7 | 73 | 57.8 | 56 | 87 | 82 | 85.6 | 91.6 | 79.4 |
| ResNet-152 | **88** | **93** | **84** | **74** | **58** | **61** | **89** | **84** | **88** | **90** | **80.9** |

Accordingly, the network performs well in IF detection but shows poor performance in detecting scapular points (LS, RS). Although most people were assumed to be healthy, they had asymmetrical protrusions in the scapula; therefore, to make the network comprehensive, this amount of difference should learn more data. Indeed, correct detection of deformities in the back surface depends on having a large dataset with different forms of spine deformities; then, the network could learn the new features. However, our algorithm has a remarkable performance, even with a limited number of subjects. As indicated in table 2, the PCK value is computed for only two networks, ResNet-50 and ResNet-152. VGG is not a suitable option for our research study because the ResNet network has been formed by deeper layers and has more computational complexities than the VGG network.

In table 2, PDJ results indicate the percentage of the points which have been detected accurately. As illustrated, ResNet-152 has better performance than ResNet-50 network, and again, the lowest level of precision is to detect the scapular points. As noted, it is because of the limited number of subjects with this specific deformity. If the network trains with various data, it can extract more complicated features. Therefore, this accuracy level of the network indicates the proper statistical population of the data.

Similarly, the following plots provide evidence for our interpretations. For example, as shown, ResNet-152 performed better than ResNet-52; the lowest accuracy was obtained for detection and estimation of spatial position in point LS. By comparing PCK and PDJ, the critical point to note is the network’s remarkable performance to detect and estimate the spatial position of the point IF. This point, placed on top of the intergluteal furrow, does not have a particular deformity compared to the other points. On the other hand, its position is not influenced by other spine deformities; therefore, enough data is fed to the network for training the features.

From the medical and clinical point of view, the manual detection of anatomical points is difficult. If the desired anatomical points are marked by several specialists on a patient’s body, none of the points are located at a specific point. Therefore, to detect these points, the spatial coordinates are not enough, and a range of neighborhoods of that point is considered. However, this research aimed to reduce the error rate of landmark detection and achieve only one coordinate as the detected point. The promising results were obtained using the proposed algorithm, and it is also essential to highlight the point that the deep convolutional network could comprehend the complexity of this type of data and meet the primary goal of research.

## 8 Discussion and Conclusion

This study aims to automatically detect the anatomical landmarks of the human back surface to investigate spinal deformities. This study’s necessity arose from the point that many people suffer from spinal deformities, among which Scoliosis is highly important because the predominant method for the detection and evaluation of the treatment process is performed through multiple radiographs. However, due to being invasive and having ionizing radiation, it increases cancer incidence, particularly among the youth. Therefore, one of the benefits of this study is the use of a non-invasive diagnosis method.

In this paper, deep learning methods were utilized since CNN has a remarkable capability, and in recent years, has been applied in a wide range of studies, obtaining remarkable results. The use of the presented algorithm is advantageous for the physician due to autonomy. Because in a spontaneous process, an appropriate diagnosis system presents recommendations to the physicians. This study’s essential attainment is deep image training because it provides rich information about the patients’ back surface to physicians.

To evaluate the algorithm’s results, the evaluation criteria were calculated and compared with ground truth. The results indicated that the network could comprehend the distinguishing features of the input and perform well in the evaluation data. As illustrated in the images, some points were not placed in the exact position, and the highest error rate in this regard was related to the scapula. This can be justified so that the network needs more data to learn more features and information.

## Notes

### Competing Interest Statement

The authors have declared no competing interest.

### Summary of Updates

The references were updated in the version

